# Stress relaxation in epithelial monolayers is controlled by actomyosin

**DOI:** 10.1101/302158

**Authors:** Nargess Khalilgharibi, Jonathan Fouchard, Nina Asadipour, Amina Yonis, Andrew Harris, Payman Mosaffa, Yasuyuki Fujita, Alexandre Kabla, Buzz Baum, José J Muñoz, Mark Miodownik, Guillaume Charras

## Abstract

Epithelial monolayers are one-cell thick tissue sheets that separate internal and external environments. As part of their function, they withstand extrinsic mechanical stresses applied at high strain rate. However, little is known about how monolayers respond to mechanical deformations. In stress relaxation tests, monolayers respond in a biphasic manner and stress dissipation is accompanied by an increase in monolayer resting length, pointing to active remodelling of cell architecture during relaxation. Consistent with this, actomyosin remodels at a rate commensurate with mechanical relaxation and governs the rate of monolayer stress relaxation – as in single cells. By contrast, junctional complexes and intermediate filaments form stable connections between cells, enabling monolayers to behave rheologically as single cells. Together, these data show actomyosin cytoskeletal dynamics govern the rheological properties of monolayers by enabling active, ATP-dependent changes in the resting length. These findings have far-reaching consequences for our understanding of developmental morphogenesis and tissue response to mechanical stress.

## Introduction

Epithelial monolayers line most of the surfaces and internal cavities of the body. They act as physical barriers that subdivide the internal body environment into discrete compartments and separate it from the external environment. To fulfil this role during embryonic development and in adult physiology, epithelia must withstand significant mechanical stresses^1-4^. During development, strain evolves slowly with strain rates of ∼0.04%.s^−1^ as the result of forces generated elsewhere in the embryo^5^; while in adult animals, strain rates of 10-100 %.s^−1^ are observed during the normal functioning of respiratory and cardiovascular systems^6-10^. In addition, organisms need to withstand external mechanical insults. Thus, for optimal tissue function and resilience, the constituent cells must be mechanically integrated to allow stresses to be spread across the whole tissue. Failure to do so can result in tissue fracture with consequences such as hemorrhage and septicemia^11-14^. Indeed, tissue fragility has been identified as a symptom in patients carrying mutations in intermediate filament and desmosomal proteins^15^, adherens junction proteins and actin cytoskeletal regulators^16-18^, and as a result of bacterial pathogens targeting intercellular adhesions^15^. At timescales of second to minutes, the ability of living tissues to dissipate stresses decreases the risks of fracture^19^, providing organisms with a protective mechanism against failure. Despite the importance of epithelial mechanics in barrier function, little is known about how epithelia dissipate stresses in response to extension.

In isolated cells, a rich phenomenology of rheological behaviours that operate at different timescales has been identified. At sub-second timescales, localised stress applied to the cell surface can be dissipated by redistribution of the fluid phase cytosol through the porous insoluble part of the cytoplasm^20^. At longer timescales, a scale-free power law rheology is observed^20,21^, which may stem from a large number of relaxation processes with different timescales operating in parallel^22^. Recent work has indicated the presence of a cut-off to the power law response imposed by the turnover rate of the actomyosin cytoskeleton^23^.

However, in tissues, the rheological behaviours observed in cells are likely to be influenced by intercellular junctions and junctional signalling^24^. Indeed, recent work has shown that adherens junctions, which link the actin cytoskeletons of adjacent cells, exhibit viscoelastic dynamics^25^. However, little is known about the stress relaxation of tissues upon deformation - despite this being an important property of many normal tissues. Nor is it known which molecular mechanisms participate in the process. In part, this derives from the difficulties of measuring stress in tissues that are mechanically coupled to a relatively thick and rigid extracellular matrix (ECM).

Here, to overcome this challenge, we study stress relaxation in epithelial monolayers devoid of an ECM subjected to a physiologically relevant strain. Our analysis reveals that at minute timescales, tissue rheology is dominated by the actomyosin cytoskeleton. Moreover, myosin contractility accelerates stress relaxation. By contrast, adherens junctions play little role in stress relaxation, acting as stable bridges connecting adjacent cells. As a consequence, the dynamics and amplitude of the relaxation of an epithelial monolayer resemble that of a single cell.

## Results

### Monolayer stress relaxation is accompanied by a change in monolayer resting length

To investigate the response of epithelia to stress, we used monolayers of Madine-Darby Canine Kidney (MDCK II) cells devoid of a substrate and suspended between test rods^11,26^. Under these conditions, all the stress in the system is borne by cells, simplifying interpretation and analysis. These monolayers were then subjected to a strain *ε*_0_ = 30% applied at a rate of 75%.s^−1^, consistent with deformations and rates observed *in vivo* under physiological conditions^7,10,27^. This 30% strain was then maintained for ∼130-140 s (**Fig 1a,b, S1**, SI), while stress relaxation was monitored. Strikingly, under these conditions, ∼70% of the stress in monolayers was found to dissipate within 60 s (**Fig 1c**). Importantly, the behaviour of monolayers was reproducible over several cycles of stress relaxation. Moreover, cells maintained their characteristic apico-basal polarity, cytoskeletal organisation, and arrangement throughout^11^.

**Figure 1:**
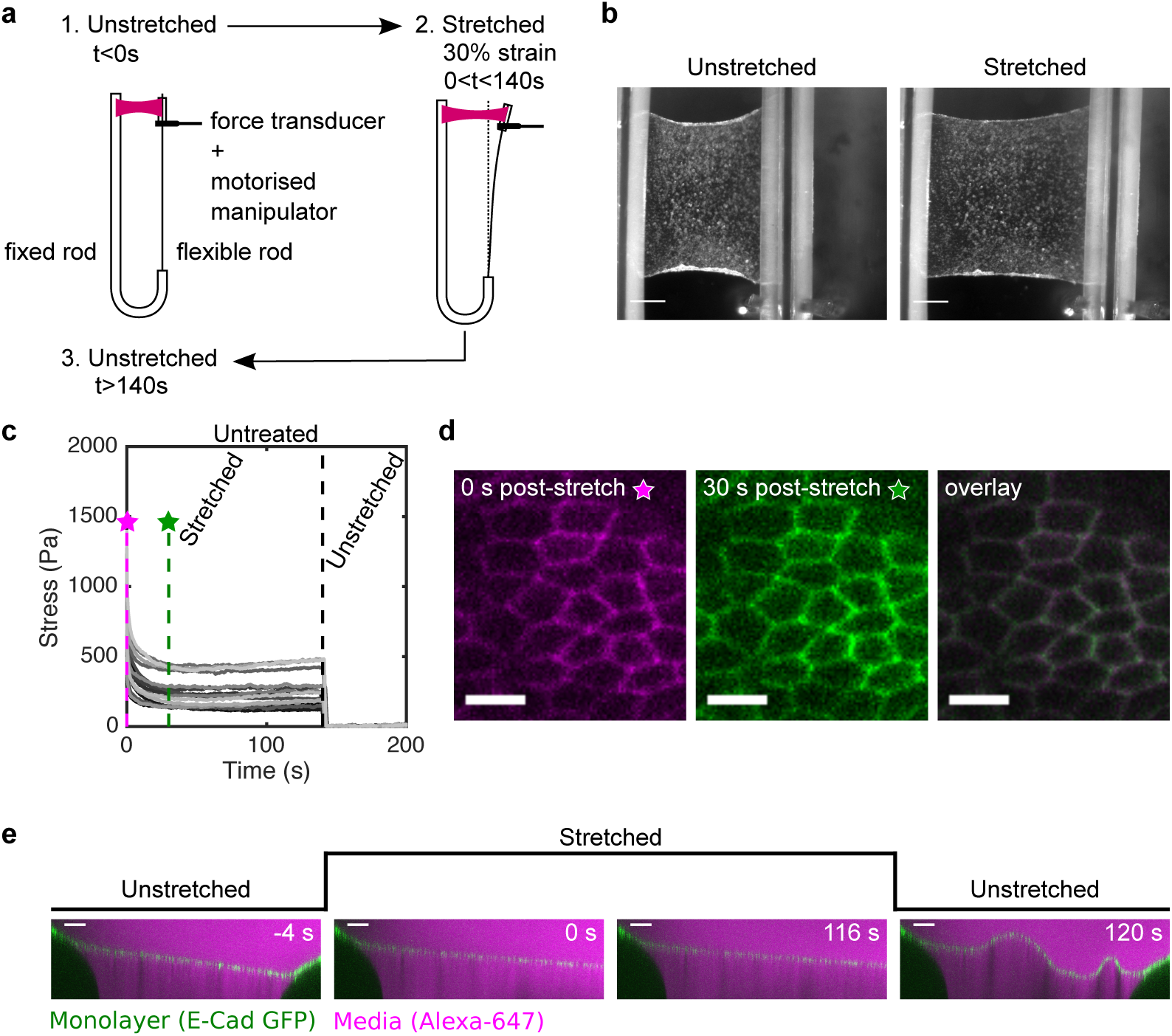
Stress relaxation in cell monolayers involves a change in resting length. (**a**) Schematic diagram of the stress relaxation experiments. Monolayers were stretched to 30% strain at a 75%.s^−1^ strain rate using a motorised micromanipulator and then kept at a fixed strain for ∼130-140 s. The flexible rod was then returned to its initial position and the monolayers were left to recover. (**b**) Bright-field microscopy images of a monolayer before and during stretch. (Scale bar: 0.5 mm) (**c**) Stress relaxation curves of cell monolayers (n=17). The magenta and green dashed lines show 0 s and 30 s after application of stretch. Stresses go to zero upon return of the flexible rod to its initial position (t=140 s, black dashed line). (**d**) Confocal microscopy images of monolayers expressing E-Cadherin GFP for 0 s (left) and 30 s (middle) after stretch. Both images were overlayed to detect potential cell shape change (right). (Scale bar: 10 μm) (**e**) Cross section of a monolayer expressing E-Cadherin GFP before application of stretch (−4 s), during stretch (0 s and 116 s) and upon release (120 s). The length of the monolayer upon release is different from its length before application of stretch. The monolayer appears in green, the surrounding medium appears in magenta due to inclusion of Alexa-647, and the test rods appear dark. (Scale bar: 100 μm)

In living tissues, stress relaxation can arise from a number of molecular- and cellular-level processes. In our experiments, however, cellular processes which typically last tens of minutes, such as oriented cell division or neighbour exchange^1,19,28^, are unlikely to contribute to stress relaxation. Indeed, when we followed cells expressing E-Cadherin GFP at high magnification, images obtained immediately after extension and 30 s later could be superimposed perfectly despite significant relaxation of stress (**Fig 1c,d**). Furthermore, cell areas and heights did not change during relaxation (**Fig S2c,d**). Together, these results indicate that stress relaxation is due to molecular-level rather than cellular-level processes.

Given the fluid-like nature of the cytoskeleton over minute to hour timescales^29^, a potential molecular origin for stress relaxation is remodelling of the cytoskeleton to adapt to the new shape of the tissue imposed by stretch. To test this hypothesis, we imaged the monolayer profile before, during, and after stretch (**Fig 1e, Video S1**).

Initially, the monolayer appears taut in between the two test rods. Application of stretch elongates the monolayer and no further changes are apparent while stretch is maintained. At the end of the experiment, however, when the test rod is returned to its initial position, the monolayer buckles. Thus, stress relaxation involves an increase in the resting length of the monolayer over time (**Fig 1e**, n=18/18 monolayers).

### Monolayer stress relaxation involves ATP-independent and ATP-dependent regimes

Next, we characterised the stress relaxation in detail. Following extension, stress relaxation began immediately. The process was biphasic, with a large amplitude fast relaxation occurring within the first ∼5 s, followed by a smaller amplitude slow relaxation, which reached a plateau after ∼60 s, as previously observed^11^ (**Fig 1c, 2a**). The presence of a plateau indicates that the material behaves like a solid at minute timescales. Examination of the relaxation curves in log-log and log-linear scales revealed that the dominant regime decays as a power law in the first phase and as an exponential in the second phase (**Fig S3**). Based on this, the relaxation can be described by a function of the form *At*^‒α^*e*^‒*t*/*τ*^ + *B* (Methods), where the kinetics of the first phase is characterised by the power law exponent *α* and the second phase by the time constant *τ*. The parameter *B*/*ε*_0_ is equivalent to an elasticity, and *A* affects the amplitude of the relaxation. Using this empirical function to describe the data, monolayer stress relaxation curves could be fit with high coefficients of determination (*r*^2^ > 0.8, n = 17 curves), without systematic bias in the residuals. The first phase had an exponent *α* = 0.28 ± 0.02 and the second phase a time constant *τ* = 11.3 ± 3.8 s. To confirm the power law nature of the first phase, we performed stress relaxation experiments for a range of deformations. After normalising the first phase to the maximum and minimum stresses, all curves collapsed onto a single master curve, consistent with scale-free rheology for the first regime (**Fig S4c-f**, SIMethods). Thus, monolayers display fluid-like properties at short timescales and solid-like properties at longer timescales. Interestingly, this behaviour was reminiscent of rheological observations in isolated cells^23,30^.

**Figure 2:**
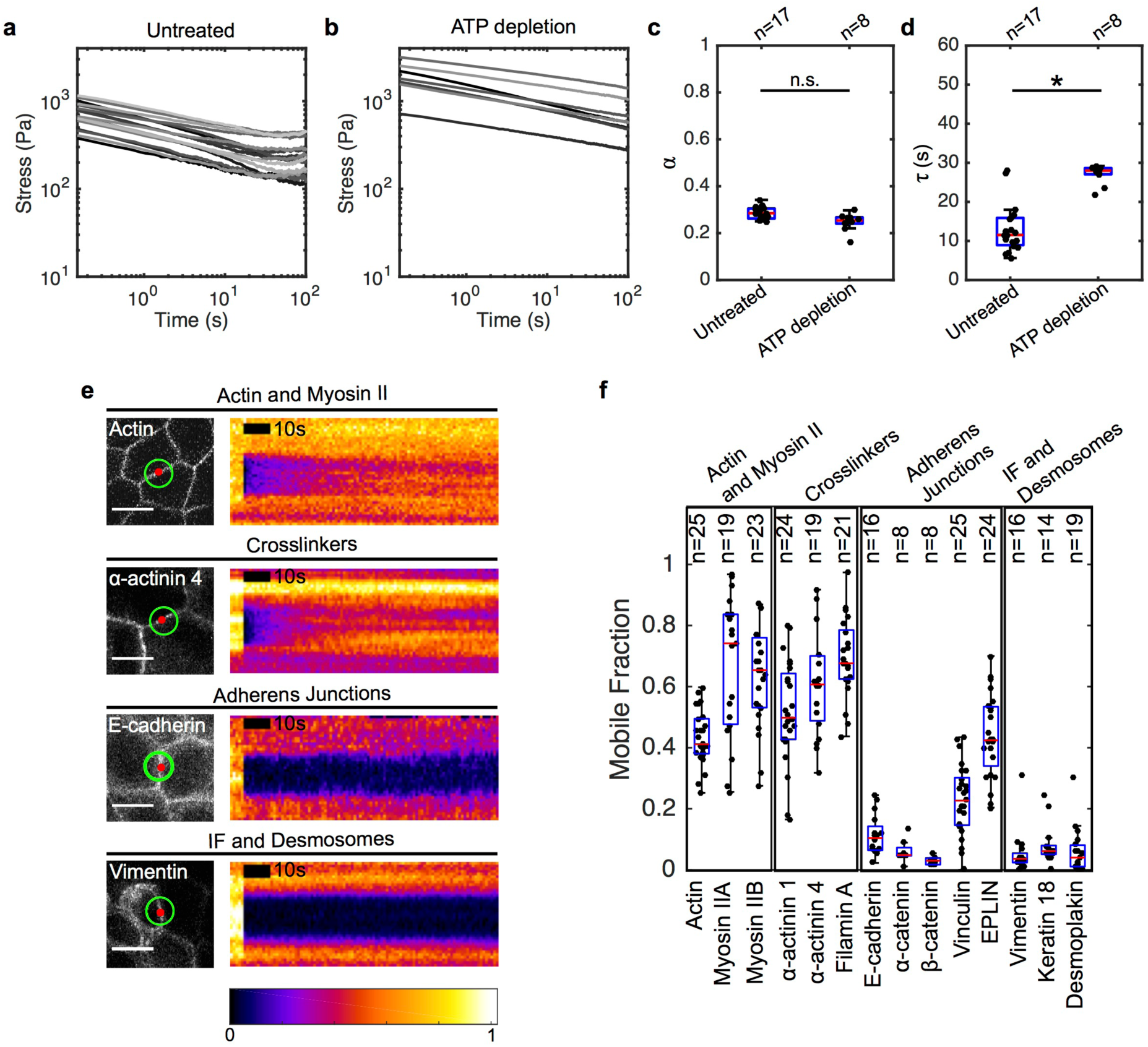
Significant cytoskeletal remodelling occurs over the timescale of stress relaxation. (**a,b**) Stress relaxation curves of untreated (**a**, n=17) and ATP depleted (**b**, n=8) monolayers plotted on a logarithmic scale. (**c,d**) Boxplots comparing the power law exponent *α* and exponential time constant *τ* of untreated and ATP-depleted monolayers. (*α*: *p* = 0.03; *τ*: *p* < 0.01) (**e**) Confocal microscopy images and kymographs of FRAP experiments. Left panels: the image shows localisation of the protein of interest, the red circle shows the bleached region, and the green circle shows the region imaged for fluorescence recovery. Right panels: each kymograph shows the normalised fluorescence intensity across the junction within the green circle. Intensities are normalised to the maximum intensity in each kymograph. (Scale bar: 10 μm) (**f**) Mobile fractions obtained from the FRAP curves for the cytoskeletal, adhesive, and junctional proteins examined.

The transition between the two phases occurs for t ∼ 5.2 s, a timescale short compared to that of biological processes involved in cell mechanics but consistent with single cell work^31^. This suggests that passive, ATP-independent processes may dominate in the power law behaviour of the first phase, while active ATP-dependent processes may dominate in the second phase. To explore this, we performed stress relaxation experiments on ATP-depleted monolayers. ATP depletion had a dramatic effect, resulting in relaxation curves with a significantly larger time constant, *τ* = 28.2 ± 0.7 s (*p* < 0.01), which appeared linear in the logarithmic scale (**Fig 2b,d**). Strikingly, however, following ATP depletion, the power law exponent was not significantly different from that observed for untreated monolayers (*α* = 0.25 ± 0.02,*p* = 0.03) (**Fig 2c**). These data suggest that the first phase of stress relaxation is ATP-independent, whereas the second phase is ATP-dependent.

### Monolayer stress relaxation depends on actomyosin not on junctional remodelling or intermediate filaments

As stress relaxation is accompanied by an increase in the resting length of the monolayer and the second phase depends upon ATP, we hypothesised that it may involve the dynamic turnover of the molecular constituents of cytoskeletal and adhesive structures. Based on previous work on the mechanics of single cells and tissues^22,32^, we decided to concentrate on the actin cytoskeleton, intermediate filaments, and the intercellular junctions that connect these structures, adherens junctions and desmosomes.

To identify the key components of each of these structures in MDCK monolayers, we used mRNA sequencing to quantify their relative abundance (SI). Although protein concentrations are not always directly correlated to mRNA transcript levels (because of differences in translation and protein degradation), they have been shown to be good predictors of protein abundance^33,34^. Moreover, low mRNA transcript levels necessarily imply low protein abundance^33,34^. Therefore, we classified proteins into categories reflecting the candidate subcellular structures (**Fig S5a**) and then selected proteins amongst the most abundant in each class for further examination.

We reasoned that only proteins that display significant turnover over the timescale of our experiments could significantly contribute to the relief of mechanical stress. To characterise turnover, we used Fluorescence Recovery After Photobleaching (FRAP) (SI). For this, we generated cell lines stably expressing GFP-tagged candidate proteins and confirmed their localisation to the relevant structures (**Fig S6**, SI). For each protein, we measured the percentage fluorescence recovery (mobile fraction) 100 s after photobleaching, because stress relaxation is complete within that timeframe. Proteins within the different candidate subcellular structures had strikingly distinct behaviours (**Fig 2e,f, S5b,c, Table 1**). Actin, myosin and crosslinkers were the most dynamic, with mobile fractions larger than 0.4 (**Fig 2f, Table 1**). In contrast, proteins of the cadherin-catenin complex, intermediate filaments and desmosomes had mobile fractions smaller than 0.1. Proteins involved in mechanotransduction exhibited intermediate levels of mobility (EPLIN and vinculin). Thus, the extent of recovery appears very different for different subcellular structures. Proteins of the actomyosin cytoskeleton (actin, myosin II and crosslinkers) have recoveries consistent with a potential role in stress relaxation. By contrast, proteins of the adherens junctions, intermediate filament networks, and desmosomes appear stable over the course of 100 s.

To confirm a potential role for actomyosin in stress relaxation, we first depolymerised F-actin using latrunculin B (**Fig 3a,b**). Loss of F-actin led to a remarkable softening of the monolayer, as seen from the 10-fold lower stresses compared to DMSO control (**Fig 2c,d**). Furthermore, relaxation curves appeared linear in the logarithmic scale, pointing to a delay in the second phase or its complete abrogation (**Fig 2d**). Together, these data show that the actin cytoskeleton underlies the active phase of relaxation and we investigated how its constituents participate in relaxation.

**Figure 3:**
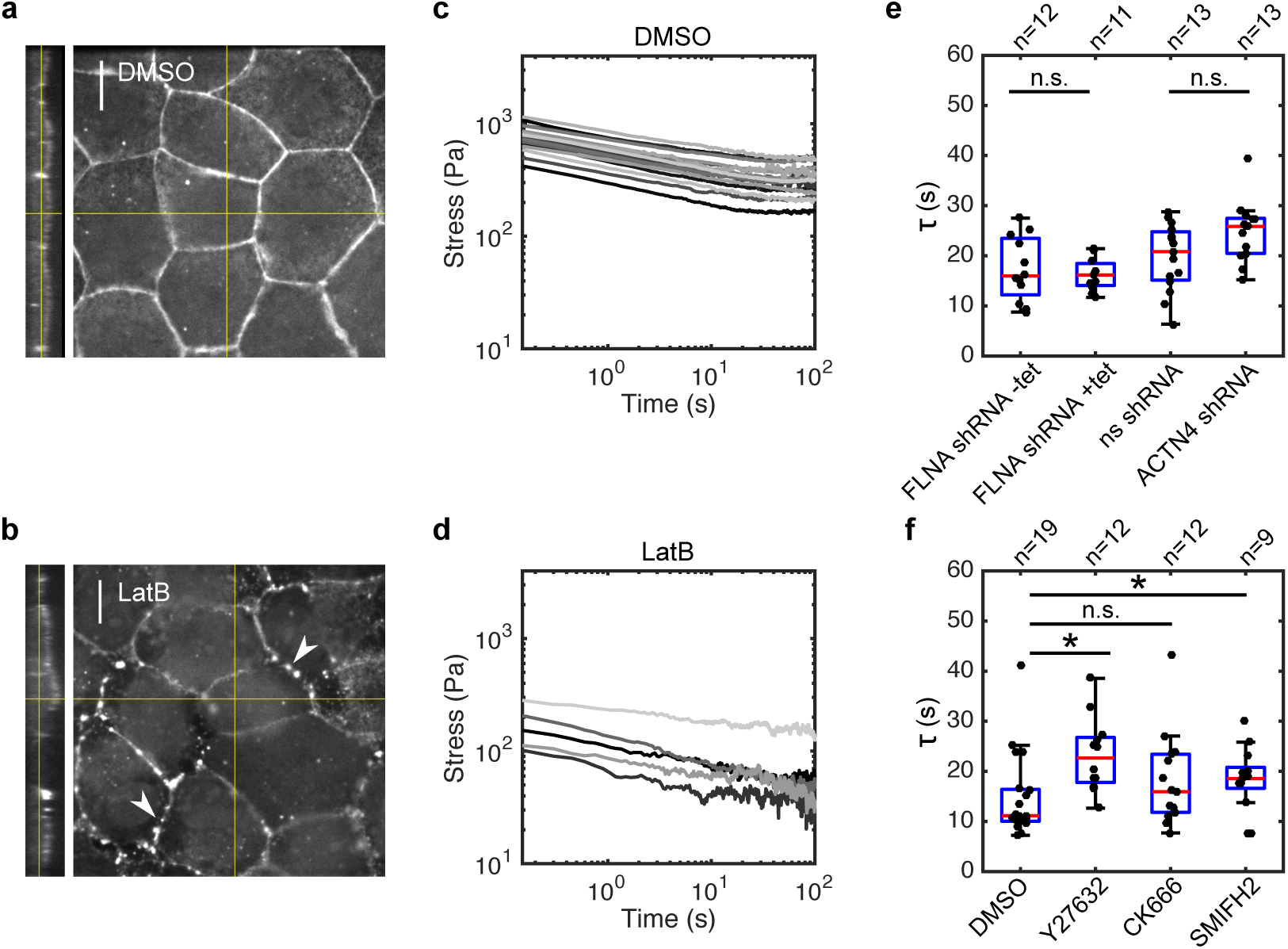
Monolayer stress relaxation is slowed by perturbations to actomyosin. (**a,b**) Confocal microscopy images showing F-actin distribution in monolayers treated with DMSO and latrunculin B for 1 h. Junctional actin localisation was perturbed following latrunculin treatment, leaving puncta of actin at the junctions (white arrows). (Scale bar: 10 μm). (**c,d**) Stress relaxation curves of monolayers treated with DMSO and latrunculin B for 1 h displayed in a logarithmic scale. (**e**) Boxplots comparing the exponential time constant *τ* in monolayers depleted for actin crosslinkers Filamin A and *α*-actinin 4. (*p* = 0.73 for FLNA shRNA +tet and *p* = 0.05 for ACTN4 shRNA, compared to their respective controls) (**f**) Boxplots comparing the exponential time constant *τ* following treatments with DMSO, Y27632, CK666 and SMIFH2 (*p* < 0.01 for Y27632 and SMIFH2 and *p* = 0.11 for CK666, all compared to DMSO).

### Active monolayer relaxation is significantly slowed by perturbing myosin contractility and actin network organisation but not crosslinkers

F-actin’s function in cytoskeletal organisation is multi-faceted: it is the basic polymer for generation of filaments, it serves as a scaffold for myosin contractility, and crosslinkers can modulate the network’s mechanics. Therefore, to examine the role of each of these functions in turn, we studied crosslinkers, myosin activity, and actin nucleation.

Crosslinkers might influence the dynamics of active relaxation by setting an intracytoskeletal friction in the actomyosin network that slows relaxation, as in single cells^23,35,36^. To investigate this, we studied stress relaxation in monolayers expressing shRNAs targeting filamin A and *α*-actinin 4, the two most abundant actin crosslinkers identified in our RNAseq experiments (**Fig S5a**). Surprisingly, the depletion of neither of the two crosslinkers had any effect on the time constant *τ* or the elasticity *B*/*ε*_0_ of monolayers (**Fig 3e, S7**). Thus, the dominant actin crosslinkers in the system do not play a role in setting the dynamics of monolayer stress relaxation.

Next, we examined the role of myosin activity. We perturbed myosin contractility using Y27632, which inhibits Rho-kinase. The treatment had a profound impact on monolayer stress relaxation. It significantly increased the relaxation time constant *τ*, leading to curves that appeared more linear in logarithmic scale (**Fig 3f, S8a**), and reduced the elasticity *B*/*ε*_0_ without affecting *A* (**Fig S8e,f**). Together, these results suggested that myosin activity accelerated the return to mechanical equilibrium following extension, as has been observed in single cells^23^.

Previous work has identified specific roles for actin networks generated through distinct nucleation pathways via the Arp2/3 complex and formins in epithelial tissues^37^. To determine the importance of actin organisation in monolayer stress relaxation, we inhibited the nucleation of actin filaments through the Arp2/3 complex using CK666, a drug that prevents Arp2/3 activation^38^, and through formins using SMIFH2, a drug that prevents barbed-end elongation via formins^39^. Arp2/3 inhibition did not significantly perturb active relaxation, whereas formin inhibition significantly increased the relaxation time constant *τ* (**Fig 3f**) – in line with formin being the dominant nucleator involved in the generation of contractile actomyosins networks in cells.

Together, these results suggest that formin-nucleated actin filaments function together with myosin II to ensure the rapid return of the monolayer to mechanical equilibrium following the application of strain.

### A phenomenological model for cell monolayer stress relaxation

Having shown that we could separate stress relaxation into a very rapid (t < 5 s) ATP-independent regime and an ATP- and actomyosin-dependent regime at longer timescales (5 < t < 75 s), we developed a simple rheological model of the system that would explain the mechanical origins of the ATP-dependent regime. In addition, because monolayer relaxation shares many commonalities with single cell relaxation, we sought to model the monolayer as an integrated mechanical system.

While cell and tissue rheology are often modelled using standard linear solid models^11^, these do not explicitly predict changes in monolayer resting length which may be important for understanding phenomena such as embryonic tissue morphogenesis, where large deformations are commonly encountered. Therefore, based on our experiments, we separated ATP-dependent monolayer mechanics into an elastic branch, which describes the response at minute long timescales using a spring *κ*, placed in parallel with an active branch, which describes the viscous transitory regime in response to mechanical perturbation (**Fig 4a**). Because of the role of myosin and changes in resting length of the monolayer during relaxation, we modelled the viscous behaviour using an active contractile element which is a spring *κ_A_* subjected to a prestrain *ε^c^*. In response to a deformation, this spring dynamically changes its resting length *L* following the equation *L̇*/*L* = *γ*(*ε^e^* – *ε^c^*), with *γ* a length-change rate, *ε^e^* = (*l_m_* – *L*)/*L* and *l_m_* the apparent length of the monolayer^40-42^. Together *κ_A_* and *ε^c^* allow the application of monolayer pre-stress *σ_c_* = *κ_A_*.*ε^c^* (Methods).

**Figure 4:**
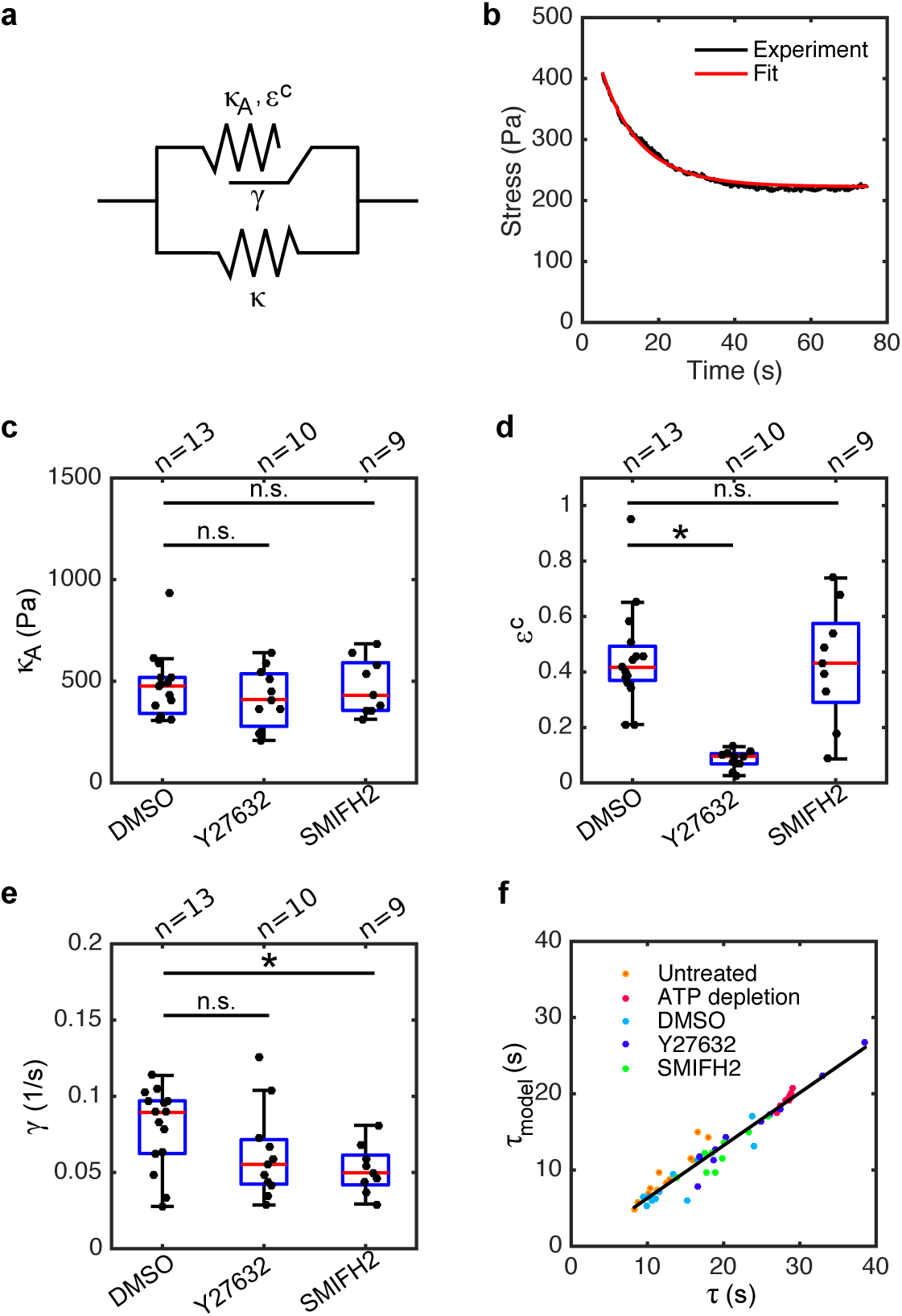
Formin-mediated actin polymerisation and myosin contractility affect different rheological properties during stress relaxation. (**a**) Diagram of the rheological model consisting of an active (top) and an elastic (bottom) branch. (**b**) The second phase of an example relaxation curve (black) is fitted with the rheological model (red). (**c,d,e**) Boxplots comparing the elastic modulus *κ_A_*, pre-strain *ε^c^* and length-change rate *γ* for monolayers treated with DMSO, Y27632 or SMIFH2. (*κ_A_*: *p* = 0.83 for Y27632 and *p* = 0.55 for SMIFH2; *ε^c^*: *p* < 0.01 for Y27632 and *p* = 0.95 for SMIFH2; *γ*: *p* = 0.02 for Y27632 and *p* < 0.01 for SMIFH2; all compared to DMSO) (**f**) Time constant *τ*_model_ calculated from the rheological model using equation (5) as a function of the time constant *τ* determined from fitting with the empirical function (l).

In experiments, the final stress in the tissue scaled linearly with strain at the timescale of minutes (**Fig S9a**), as expected for solid, spring-like behaviour. Consistent with the presence of an active element, we confirmed the presence of pre-stress in monolayers prior to extension with values *σ_c_*∼130 Pa (SI, **Fig S9b**). We then modelled the actomyosin-dependent stress relaxation in the system using *κ_A_*,*γ*, and *ε^c^*, yielding a characteristic time *τ*_model_∼1/[*γ*(1 + *ε^c^*)]. Next, we fitted the second phase of stress relaxation using the analytical solution to determine *κ_A_* and *γ*, knowing *σ_c_* = *κ_A_*.*ε^c^* (**Fig 4b**). Analytical curves fitted the experimental data well (*r*^2^ > 0.8 for 80% of the relaxation curves) without any systematic bias in the residuals, yielding *κ_A_*∼142 Pa and *γ*∼0.09 s^−1^. Thus, by introducing an active element that adjusts the monolayer resting length, we were able to reproduce monolayer stress relaxation in the ATP-dependent phase.

### Myosin contractility contributes to stress relaxation by generating pre-strain and crosslinking actin filaments

To understand the link between mechanical behaviour and biological mechanisms, we analysed perturbation experiments using our rheological model as a guide. To do this, we measured changes to *κ* and *σ_c_* from our experimental data and obtained values for *κ_A_*, *γ*, and *ε^c^* from curve fitting with the condition *σ_c_* = *κ_A_*.*ε^C^*. This revealed different action mechanisms for formins and myosin II contractility. Rho-kinase inhibition led to a decrease in *κ*, a five-fold reduction of pre-strain *ε^c^*, a partial decrease in length-change rate *γ* (*p* = 0.02), but did not affect the stiffness *κ_A_* of the active element. Thus myosin contributes to both the viscous and the elastic branches of the model, perhaps through its contractility and crosslinking functions respectively. By contrast, the inhibition of formin activity only altered the length-change rate *γ* (**Fig 4c-e, S9c**). ATP depletion led to a two-fold increase in *κ_A_* and to a significant decrease in both *ε^c^* and *γ* (**Fig S10a-c**). Importantly, the characteristic time constants *τ*_model_ computed from the model correlated well with those determined from empirical fitting (**Fig 4f**), validating our approach.

Together, these results indicate that the pre-strain *ε^c^* depends on contractility alone, *γ* depends on actin polymerisation, and *κ* depends on actin organisation and density.

## Discussion

Here, we characterise stress relaxation in monolayers and the molecular turnover of the stress-bearing biological structures. Our data paint a picture in which intercellular junctions form stable interconnections between cells allowing the monolayer to behave as a single cell with its rheology controlled by actomyosin. Together F-actin remodelling and myosin contractility endow the monolayer with solid-like mechanical properties at minute timescales, act as driving forces to reach a new mechanical steady-state following extension, and regulate the resting length of the monolayer.

### Monolayer rheology is governed by actomyosin at second to minute timescales

When we examined the response of suspended epithelial monolayers to a step deformation, we found that ∼70% of stress is dissipated within 60 s and that relaxation can be described by a power law with an exponential cut-off at longer timescales. Examination of the temporal evolution of cell morphology revealed that dissipation occurred through molecular-rather than cellular-scale processes. Interestingly, the two phases of relaxation were also distinguished by their dependence upon ATP.

The first power law phase did not depend upon ATP and had an exponent *α* ∼ 0.28, similar to that reported for single cells subjected to a step extension in a geometry similar to our experiments^43-45^ and for cell aggregates subjected to compression^46^. Given the dependence of the second phase of relaxation on ATP, we focused on the contributions of subcellular structures known to play a role in cell and tissue mechanics such as the adherens junctions^11,47,48^, desmosomes^49^, intermediate filaments^50,51^ and actomyosin^11,52-56^. Stress relaxation within cytoskeletal and adhesive structures likely stems from molecular turnover of their constituents^57^. In support of this idea, we found a clear separation in the extent of turnover of these structures within the 100 s duration of our experiments. Intermediate filaments and desmosomes display very little turnover within the 100 s duration of our experiments^58-60^ (**Fig 2**, **S5**), suggesting that they do not significantly participate in stress relaxation. Proteins of the cadherin-catenin adhesive complexes also display little turnover over the timescale of stress relaxation^61^, whereas proteins of the actomyosin cytoskeleton turn over significantly (**Fig 2**, **S5**). Furthermore, we found that treatments that target actomyosin lead to significant changes in stress relaxation (**Fig 3, S8**), suggesting that actomyosin governs monolayer rheology at second to minute timescales.

### Monolayer rheology is similar to single cell rheology

Interestingly, in line with actomyosin controlling rheology, stress relaxation in monolayers displayed many similarities to stress relaxation in single cells. This is surprising, since single cells lack adhesive structures. However, when subjected to 25-30% extension, both single cells and monolayers displayed an initial phase of relaxation following a power law with a similar exponent before reaching a plateau at longer timescales corresponding to ∼20-30% of the initial stress^30,43^. The existence of such a plateau indicates that both single cells and monolayers switch from a liquid-like behaviour at short timescales to a solid like behaviour on minute-long timescales. This switch may represent the passage from a regime dominated by cytoplasmic rheology to one dominated by actomyosin at the cell periphery^20,41^.

In our experiments, stress relaxation was well-described by a power law with an exponential cut-off, something that has also been observed in experiments probing the rheology of the cortex of single-cells^23^. The time constant *τ* of the exponential cut-off was similar for single cells and monolayers (∼10 s) and inhibition of myosin contractility led to a two-fold increase in *τ* and significantly slower relaxations. Intriguingly, our mechanical model suggested that myosin participated in monolayer mechanics both through its contractility affecting *ε^c^* and through a crosslinking role affecting *κ* (**Fig 4**, **S9**). Why crosslinking via myosins dominates over crosslinking by specialised proteins is unclear (**Fig 3**). One possibility is that myosin mini-filaments themselves may have long lifespans providing stable crosslinking. In this picture, although individual myosins within a mini-filament are progressively replaced, the mini-filament retains its position within the network. Myosins may thus contribute to network elasticity at minute timescales, while specialised crosslinkers contribute to the viscous behavior of the monolayer.

Our analysis also explains why monolayers behave like single cells. Our measurements of protein turnover show that adhesive structures turn over significantly less than actomyosin. One consequence of these clear differences in turnover extent is that adherens junctions form stable interconnections between cells allowing the monolayer to behave as a single cell with its rheology controlled by actomyosin. This further implies that multicellular rheology in monolayers may be controlled by emergent properties of actomyosin gels at the molecular-level^29,62^, something that will form an interesting direction for future work.

### Myosin contractility and F-actin remodelling control stress relaxation through separate pathways

Theoretical predictions and experiments suggest that actomyosin rheology is controlled by crosslinker dynamics, myosin contractility, and actin polymerisation dynamics^63-65^. Depletion of the two most abundant F-actin crosslinkers (*α*-actinin 4 and filamin A) did not perturb the ATP dependent phase of relaxation (**Fig 3**), in contrast to what is observed in single cells^23^, perhaps because of differences in protein abundance or organisation due to the formation of intercellular junctions. Surprisingly, we found that blocking myosin contractility and inhibiting formin-mediated actin polymerisation both significantly increased the duration of relaxation (**Fig 3**). To distinguish the mechanisms of these two perturbations, we used an equivalent rheological model in which active and passive parts are explicitly separated. Importantly, we also allow the monolayer resting length to change by introducing an active element that responds to monolayer deformation by adapting resting length with a certain timescale. Analysis of perturbation experiments showed that inhibition of contractility only affects pre-strain; whereas formin inhibition affects length-change rate only. Within our model and our experiments, both lead to slower return to mechanical steady-state following a step extension but do not prevent relaxation. Thus, myosin contractility and formin-mediated actin polymerisation both accelerate the return of monolayers towards mechanical steady state following extension. Although our model can replicate our experimental data faithfully, further work will be necessary to determine its predictive power.

How stress is relaxed in cells and monolayers remains poorly understood. Previous theoretical and experimental studies have suggested that changes in the resting length of cells and tissues may underlie stress relaxation^40,41^. In line with this, we showed that monolayer resting length increases in response to a sustained stretch (**Fig 1**). This length change appears to depend on formin-mediated polymerisation, although the detailed molecular mechanism remains to be determined (**Fig 4**). Increase in monolayer resting length must originate from changes in the resting length of its constituent cells, thus our experiments suggest that stress relaxation in single cells also involves cytoskeletal remodelling. The realisation that monolayers can change resting length in response to stress may have important consequences for our understanding of developmental morphogenesis. Indeed, many developmental processes involve large tissue deformations in response to stress generated elsewhere in the embryo. Therefore, changes in monolayer resting length may participate to relax stress in the tissue alongside cellular-level processes, such as neighbour exchanges or oriented division.

In summary, our data paint a picture in which actomyosin plays a central role in monolayer mechanics endowing the monolayer with solid-like mechanical properties at minute timescales, acting as a driving force to reach a new mechanical steady state and regulating monolayer resting length. The challenge will now be to understand which properties of actomyosin gels at the molecular-level control tissue-scale mechanics.

## Code availability

Custom-written code used for data analysis are available from the authors upon request.

## Data availability

All data supporting the conclusions are available from the authors on reasonable request.

## Acknowledgements

The authors wish to acknowledge present and past members of the Charras, Baum, Kabla, and Muñoz labs for stimulating discussions. The authors acknowledge technical support from UCL Genomics for sequencing and analyzing total RNA data. N.K. was funded by the Rosetrees Trust, the UCL Graduate School, the EPSRC funded doctoral training program CoMPLEX, and the European Research Council. N.K. was in receipt of a UCL Overseas Research Scholarship. N.K. was supported by the Prof Rob Seymour Travel Bursary Fund for research visits to Barcelona. J.F. is funded by BBSRC grant (BB/M003280 and BB/M002578) to G.C. and A.K.. G.C. is supported by a consolidator grant from the European Research Council (MolCellTissMech, agreement 647186). J.J.M, N.A. and P.M. acknowledge the support of the Ministry of Economy, Industry and Competitiveness (MINECO) through Grants No. DPI2013-43727R, and DPI2016-74929-R and the Generalitat de Catalunya through Grant No. 2014-SGR-1471. N.A. is also financially supported by Universitat Politècnica de Catalunya (UPC) and Consorci Escola Industrial de Barcelona (CEIB) through Grant UPC-FPI 2012, and the European Research Council under the European Community’s 7th Framework Programme (FP7/2007-2013)/ERC Grant Agreement No. 240487. P.M. is also supported by the European Molecular and Biology Organisation (EMBO) under grant ASTF 351-2016. B.B. was supported by UCL, a BBSRC project grant (BB/K009001/1) and a CRUK programme grant (17343). A.K. was supported by BBSRC grants (BB/K018175/1, BB/M003280 and BB/M002578). M.M. was supported by EPSRC (EP/K038656/1). A.Y. was supported by an HFSP Young Investigator award to G.C. (RGY 66/2013).

## Author contributions

N.K., A.H. and G.C. designed the experimental setup. N.K., A.K., B.B. and G.C. designed the experiments. N.K. carried out the relaxation experiments. G.C. carried out FRAP experiments and protein localisation experiments. A.Y. carried out western blot experiments. N.K. carried out most of the data and image analysis. J.F. carried out image analysis to measure pre-stress. A.K. contributed to theoretical analysis. N.A., P.M. and J.J.M. designed the rheological model. J.J.M contributed to computational analysis. A.K., J.J.M and M.M. provided conceptual advice. Y.F. provided cell lines. N.K., B.B. and G.C. wrote the manuscript. All authors discussed the results and manuscript.

## Methods

### Cell culture and generation of cell lines

MDCK II cells were cultured at 37°C in an atmosphere of 5% CO2 in air in high glucose DMEM (ThermoFisher) supplemented with 10% FBS (Sigma) and 1% penicillin-streptomycin (ThermoFisher). Mechanical experiments and imaging were performed in Leibovitz’s L15 without phenol red (ThermoFisher) supplemented with 10% FBS.

In order to visualise the junctional and cytoskeletal structures, as well as to determine the turnover kinetics of various proteins, stable lines of MDCK II cells expressing the following proteins were used: E-Cadherin GFP, actin GFP, *α*-catenin GFP, *β*-catenin GFP, vinculin GFP, EPLIN GFP, *α*-actinin 1 GFP, *α*-actinin 4 GFP, filamin A GFP, vimentin GFP, keratin 18 GFP, desmoplakin GFP, NMHCIIA GFP and NMHCIIB GFP.

Cell lines expressing E-Cadherin GFP and keratin 18 GFP were described in Harris et al.^11^. Other cell lines were generated by linearisation of plasmids encoding the FP tagged protein of interest with the appropriate restriction enzyme. The following plasmids were used: *α*-catenin GFP (a kind gift of Dr E Sahai, the Francis Crick Institute, UK), *β*-catenin GFP (a kind gift of Dr Beric Henderson, University of Sydney, Australia), vinculin GFP (a kind gift of Prof Susan Craig, Johns Hopkins University, USA), EPLIN GFP (a kind gift of Prof Elizabeth Luna, University of Massachusetts, USA, Addgene plasmid 40947), *α*-actinin 1 GFP^66^, *α*-actinin 4 GFP (a kind gift of Prof Doug Robinson, Johns Hopkins University, USA), filamin A GFP (a kind gift of Dr Paul Shore, University of Manchester, UK), vimentin GFP (a kind gift of Prof Robert Goldman, Northwestern University, USA), desmoplakin GFP (a kind gift of Prof Kathleen Green, Northwestern University, USA, Addgene plasmid 32227), NMHCIIA GFP and NMHCIIB GFP (both kind gifts of Dr Robert Adelstein, National Heart, Lung and Blood Institute, USA, Addgene plasmids 11347 and 11348). The cell line expressing actin GFP was generated by inserting actin-GFP into a retroviral vector (pLPCX, Takara Clontech), generating retrovirus as described in Harris et al.^11^, and transducing it into MDCK cells. To create all other stable cell lines, the plasmid of interest was first linearised with the appropriate restriction enzyme and then transfected into wild type MDCK II cells using electroporation (Lonza CLB). ∼10^6^ cells were transfected with 10 μg (NMHCIIA-GFP, NMHCIIB-GFP) or 2 μg (all other plasmids) of cDNA according to manufacturer’s instructions and then selected with antibiotics for 2 weeks. In order to achieve a homogenous level of fluorescence expression, cells were sorted using flow cytometry. Cells expressing E-Cadherin GFP were cultured in presence of 250 ng.ml^−1^ puromycin. Cells expressing actin GFP were selected in presence of 1 μg.ml^−1^ puromycin. All other cell lines were selected in presence of 1 mg.ml^−1^ G418.

To study the role of crosslinkers, cell lines stably expressing shRNA targeting filamin A and *α*-actinin 4 were used. Filamin A shRNA was expressed in a tetracycline-inducible manner^67^. These cells were cultured in presence of 5 μg.ml^−1^ blasticidin and 800 μg.ml^−1^ G418. To induce expression of shRNA, cells were incubated in presence of 2 μg.ml^−1^ doxycycline for 72 h prior to the experiments. Plasmids of encoding non-silencing shRNA and shRNA targeting *α*-actinin 4 were a kind gift from Prof Bill Brieher (University of Illinois, USA). Following linearisation of the plasmids, stable cell lines expressing control shRNA and *α*-actinin 4 shRNA were generated by transfecting the plasmids into wild type cells using electroporation (Lonza CLB) as described above. Control and *α*-actinin shRNA lines were amplified and selected in presence of 4 μg.ml^−1^ puromycin. Protein depletion was ascertained using Western blotting.

### Generating suspended cell monolayers

Suspended cell monolayers were generated as described by Harris et al.^11,26^. Further information is provided in SI.

### Mechanical testing procedure

The mechanical testing setup was assembled on top of an inverted microscope (Olympus IX-71) (**Fig S1a**). First, the petri dish containing the stress measurement device was secured on the microscope stage with 4 pieces of plasticine. The force transducer (SI-KG7A, World Precision Instruments) with a tweezer-shaped mounting hook (SI-TM5-KG7A-97902, World Precision Instruments) was mounted on a 3D motorised micromanipulator (Physik Instrumente) with a custom-made adaptor. The fixed rod of the device was held with the arm of a 3D manual micromanipulator, while the top Tygon section of the flexible rod was held with the tip of the force transducer. Both motorised and manual micromanipulators were equipped with a magnetic plate that secured them to the custom-made metal stage of the microscope.

Using the motorised micromanipulator, the monolayers could be extended to different strains with controlled strain rates. Extended monolayers exerted restoring forces on the flexible rod, causing the transducer tip to bend. The extent of bending was translated into a voltage value that was converted into a digital signal using a data acquisition system (USB-1608G, Measurement Computing) and recorded onto a computer. Both the data acquisition system and the motorised micromanipulator were controlled with a custom-written code in Labview. The monolayer and the transducer tip were imaged every 0.5 s using a 2× objective (2× PLN, Olympus).

The mechanical testing procedure consisted of several steps:

- Initial approach: The tip of the force transducer was initially brought into contact with the Tygon tubing and then positioned such that the left tweezer arm was out of contact but within 50 μm distance from the Tygon tubing. This enabled identification of the contact point of the transducer tip with the device during the mechanical testing procedure.
- Preconditioning: The monolayers were subjected to 8 cycles of loading to a 30% target strain at a 1%.s^−1^ strain rate. This ensured breakage of any residual collagen attached to the monolayer (especially close to the rods), as well as causing the samples to evolve into a “preconditioned” state, where the slope of the stress-strain curve did not change in successive cycles, and hence, several experiments could be conducted on the same sample with a high degree of reproducibility.
- Stress relaxation experiments: The monolayers were extended to 30% strain at a 75%.s^−1^ strain rate and then kept at a fixed 30% strain for ∼130-140 s. The micromanipulator was then returned to its initial position before stretch. This released the monolayers and they were left unstretched for ∼130-140 s to recover to their initial state. This stress relaxation experiment was repeated 3 times on each monolayer.
- Loading until failure: The monolayers were extended until failure at 1%.s^−1^ strain rate. After rupturing the monolayer, the flexible rod was returned to its initial position.
- Calibration of the device: To allow conversion from voltages to force, the device was calibrated. For this, the wire was extended at the same rate and to the same extent as in the cycling experiments. This was repeated 5 times. The length of the wire *L_w_* was measured using a Canon FD macro-lens (Canon, Surrey, UK) interfaced to a Hamamatsu EMCCD camera (Hamamatsu Orca ER, Hamamatsu UK, Hertfordshire, UK) (**Fig S1b**).

### Analysis of the relaxation curves

To analyse the response of monolayers to a step deformation, the first 75 s of the stress relaxation curves were fitted with a function comprising a power law with an exponential cut-off:

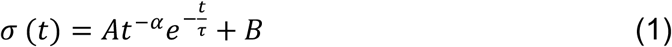

The fitting procedure was as follows: First, the initial conditions for the fitting were determined. *B* was the residual stress after the curves plateaued and was defined as the average stress for 70 s < t < 75 s. *A* + *B* was defined as the initial stress at the second timepoint (t = 0.150 s) after the step deformation. The first timepoint after application of the step deformation was ignored in order to allow the calculations to be performed in the logarithmic scale (**Fig S3**). To estimate *α*, the first 5 s of the curves were used. In practice, *σ*(*t*<5 *s*)–*B* was plotted as a function of time in the logarithmic scale and fitted with a line, with *α* being the slope of this line. To estimate *τ*, *σ*(5 < t < 20 s) – *B* was plotted in a semi-logarithmic scale and fitted with a line, with *τ* being the slope of this line (**Fig S3**). For each experimental measurement, *B* was kept constant and the relaxation curve was fitted using equation (1), with the free parameters *A*, *α* and α. The trust-region-reflective least squares algorithm, a built-in Matlab fitting procedure, was used for the fitting. The fitting was performed for the three individual repeats of the stress relaxation experiments on each monolayer. The fitted values obtained from the three repeats were then averaged to obtain a single value for each parameter.

The goodness of fit was determined using the coefficient of determination *r*^2^ and curves with *r*^2^ < 0.80 were excluded from further analysis. This represented less than 4% of experimental curves acquired. Outliers were determined as described in the statistical analysis section and the curves for which either of the three fitted parameters *A*, *α* and *τ* were outliers were not included for statistical analysis. On average, less than 10% of the data was excluded from analysis.

### Confocal microscopy for cell area and height measurements

High magnification devices were prepared as described in SI High magnification imaging devices. Images for cell area measurements were obtained using either a scanning laser confocal microscope (Olympus IX-81 with an FV-1000 confocal head) with a 20× objective (UPLSAPO, Olympus, N.A.=0.75, working distance: 0.6 mm) or a spinning disk confocal microscope (Yokogawa) with a 40× objective (UPLSAPO, Olympus, N.A.=0.9, working distance: 0.18 mm). Images for cell height measurements were obtained using a scanning laser confocal microscope (Olympus IX-81 with an FV-1000 confocal head) with a 30× silicone oil immersion objective (UPLSAPO, Olympus, N.A.=1.05, working distance: 0.8 mm).

### Confocal microscopy for protein localisation

Cells were imaged using either a scanning laser confocal microscope (Olympus IX81 with a FV1000 confocal head or Olympus IX83 with a FV1200 confocal head) or a spinning disk confocal microscope (Yokogawa). Images were taken using a 100× oil immersion objective (UPLSAPO, Olympus, N.A.=1.4, working distance: 0.13 mm) and confocal stacks were acquired at 0.3 μm intervals in z.

### Fluorescence Recovery After Photobleaching

Fluorescence Recovery After Photobleaching (FRAP) experiments were performed using a 100× oil immersion objective on a scanning laser microscope. The protocol for the FRAP was as described by Fritzsche et al.^68^. Further information is provided in SI.

### Chemical treatments

To deplete monolayers from their ATP stocks, the high glucose growth medium was gradually exchanged with PBS and the monolayers were gently washed with PBS to ensure that no high glucose medium remained on the monolayer and in the dish. PBS was then replaced with a solution of sodium azide (4 mM) and 2-deoxyglucose (10 mM) in imaging medium. After 45-60 min incubation at 37°C, collagenase type-II was added to the medium to enzymatically remove the collagen. The monolayers were then incubated at room temperature for 1 h. Finally, the medium was exchanged for imaging medium containing sodium azide and 2-deoxyglucose.

To depolymerise F-actin, monolayers were treated with 3 μM Latrunculin B (Calbiochem). To inhibit polymerisation of F-actin through formins or Arp2/3 complex, monolayers were treated with 40 μM SMIFH2 (Calbiochem) or 100 μM CK666 (Calbiochem), respectively^38,39^. After digestion of collagen, the medium was gradually replaced with the imaging medium. The drugs were then added to the imaging media and the monolayers were incubated with the drug for 1 h at 37°C. The experiments were performed in presence of the drugs.

For inhibiting myosin contractility, monolayers were treated with 50 μM Y27632 (Calbiochem). After digestion of collagen, the medium was gradually replaced with the imaging medium. Finally, the relevant dose of Y27632 was added just before starting the experiments. Control experiments were carried out in the presence of DMSO alone.

### Immunostaining

Cells were incubated with Leibovitz’s L15 for 5 min at room temperature, before being fixed with 4% PFA diluted in L15 at room temperature for 15 min. After three 10 min washes with PBS, cells were permeabilised with 0.5% Triton X in PBS for 5 min on ice. To block nonspecific binding, cells were incubated in 10 mg.ml^−1^ BSA in PBS for 10 min at room temperature and then washed 3 times with BSA/PBS, with each wash lasting 10 minutes. Next, cells were incubated with Phalloidin 647 (Life technologies, A22287, 1:200 dilution) for 1 h at room temperature. Finally, cells were washed three times with BSA/PBS for 10 min each before being mounted in Fluorsave (Merck Millipore) for imaging.

### Fitting the second phase of the relaxation with the rheological model

The second phase of the relaxation curves (defined for t > 5.2 s) was fitted with the rheological model shown in **Fig 4a**. The resting length *L* of the active branch has the following evolution law:

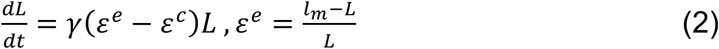

where *γ* is the rest-change rate, *ε^c^* is the pre-strain and *l_m_* is the apparent length of the monolayer. Following application of a step strain at t = 0 s from *l_m_* = *L*_0_ to *l_m_* = *L*_1_, the resting length will evolve. Since the monolayers are pre-stressed and contractile, the initial value of the resting length is given by *L*(0) = *L*_0_/(1 + *ε^c^*). This provides the initial pre-strain: *ε^c^* = [*L*_0_ - *L*(0)]/*L*(0). Knowing that *σ* = *κ_A_ε^e^*, this will lead to stress relaxation of the form:

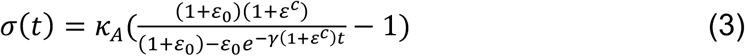

where *κ_A_* is the spring stiffness and *ε*_0_ is the applied strain defined as 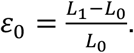 The characteristic time *τ*_model_ for this relaxation can be calculated as:

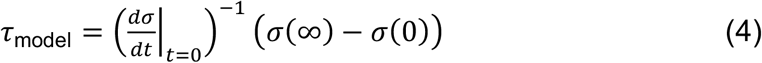

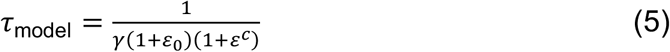

The fitting was performed as follows: first the residual stress *B* was subtracted from the total stress. Knowing that the measured pre-stress *σ_c_* is equal to *κ_A_*.*ε^C^*, the remaining stress was fitted with the stress relaxation function given in equation (3), allowing *κ_A_* and *γ* to vary. The goodness of fit was determined using the coefficient of determination *r*^2^ and curves with *r*^2^ < 0.80 were excluded from further analysis. This represented less than 15% of the analysed curves. Outliers were determined as described in the statistical analysis section and the curves for which either of the three fitted parameters *κ_A_*, *γ* or *ε^c^* were outliers were not included for statistical analysis. On average, ∼20% of the data was excluded from analysis.

### Statistical analysis

All data analysis and curve fitting were conducted using custom-written code in Matlab. For each dataset, outliers were defined as the values that fell outside the range [*q*_1_ – *w* × (*q*_3_ – *q*_1_),*q*_3_ +*w* × (*q*_3_ – *q*_1_)], where *q*_1_ and *q*_3_ were the 25th and 75th percentiles of the data and *w* was 1.5. Outliers were excluded from statistical analysis. The normality of the data was tested using both Lilliefors and Shapiro-Wilk tests in R, which confirmed non-normality of some datasets. Statistical analysis was performed in Matlab, using a Wilcoxon rank sum test that does not assume normality of the data. Datasets with *p* < 0.01 were deemed to be significantly different. For all boxplots, the edges of the box represent the 25th and 75th percentiles of the data, the red line marks the median and the whiskers extend to include the most extreme data points that are not considered to be outliers. Points on each boxplot represent individual monolayers or cells. Each dataset is pooled across experiments performed on at least 3 individual days.

